# Iron promotes Slc7a11-deficient valvular interstitial cell osteogenic differentiation: A possible mechanism by which ferroptosis participates in intraleaflet hemorrhage-induced calcification

**DOI:** 10.1101/2021.09.06.459126

**Authors:** Ran Xu, Dan Zhu, Jianghong Guo, Ying Huang

**Author notes:** Ran Xu and Chong Wang contributed equally to this study. Danzhu and Jianghong Guo are the co-corresponding authors of this study.

## Abstract

**Background:** Calcific aortic valve disease (CAVD) is the most frequent pathogeny of aortic valve replacement in developed countries. Iron deposits are found in the intraleaflet hemorrhage (IH) areas of calcific aortic valves. Ferroptosis is a form of regulated cell death that involves metabolic dysfunction resulting from iron overload-dependent excessive lipid peroxidation. In this research, we attempted to clarify the role of ferroptosis in CAVD.

**Methods:** The level of ferroptosis in tissue and valvular interstitial cells (VICs) was assessed by the contents of 4-HNE, NADPH, ROS, and GSH, lipid peroxidation and mitochondrial morphology. The levels of calcification, iron accumulation and Slc7a11 expression in surgical aortic valve specimens were detected by Alizarin red or Von Kossa, Perl’s blue and immunohistochemical staining. The osteogenic differentiation of VICs was assessed by PCR and western blot analyses. Furthermore, RNA sequencing was used to detect potential differentially expressed genes between normal and osteogenic medium-treated (OM-treated) VICs.

**Results:** Our experiments demonstrated that ferroptosis occurred in the IH areas of calcific aortic valves. We also found that Slc7a11 was expressed at low levels in OM-treated VICs and IH areas. Finally, we demonstrated that iron promoted Slc7a11-deficient VICs osteogenic differentiation by aggravating ferroptosis in vitro.

**Conclusion:** In conclusion, iron promotes Slc7a11-deficient VIC osteogenic differentiation by aggravating ferroptosis in vitro, thereby accelerating the progression of aortic valve calcification.

**Statement:** Statements and opinions expressed in the articles and communications herein are those of the author(s) and not necessarily those of the Editor(s), Society, or publisher, and the Editor(s), Society, and publisher disclaim any responsibility or liability for such material.

**Brief Summary:** In this work, two novel notions have been proposed. First, we reported that ferroptosis participated in the progression of CAVD. Second, this is the first cytology experiment of valvular interstitial cells (VICs) to clarify the mechanism by which intraleaflet hemorrhage aggravates calve calcification. This research provides new ideas and targets for alleviating the progression of CAVD, especially in patients who have calcified aortic valves without severe stenosis.

## Introduction

Calcific aortic valve disease (CAVD) is the most frequent pathogeny of surgical aortic valve replacement in developed countries[1]. As a population ages, the incidence of CAVD continuously increases[2]. However, no approved pharmacological treatments are available to stop or alleviate the progression of CAVD. Therefore, it is necessary to elucidate the mechanisms of calcification to ultimately develop alternative prophylaxis and therapeutic strategies for CAVD.

Intraleaflet hemorrhage (IH) is common in degenerative aortic valve disease[3]. Previous studies have demonstrated that IH is associated with the rapid progression of degenerative aortic valve stenosis[4–6]. However, these studies lacked related cellular experiments, and the mechanisms underlying these pathological processes still need to be explored. In the IH area, iron accumulates as a result of neovascularization or heme metabolism in macrophages[5,6]. Recently, increasing histological evidence has suggested that iron may be associated with calcification in CAVD[5,7]. However, detailed mechanism studies on how iron overload accelerates the pathological progression of CAVD are still lacking.

Ferroptosis is a form of regulated cell death that involves metabolic dysfunction resulting from iron overload-dependent excessive lipid peroxidation. Many studies showed iron accumulation in IH areas. Our previous research also reported that IL-18 promotes erythrophagocytosis and erythrocyte degradation to aggravate iron overload[8]. Moreover, oxidative stress induces ferroptosis[9], which also participates in the progression of CAVD[1]. All of these findings led us to explore the relationship between ferroptosis and valve calcification.

In this research, we explored the specific mechanism of differentiated VICs in iron overload-induced calcification. We also attempted to clarify the role of ferroptosis in the development of aortic valve calcification. Our results suggest that IH and ferroptosis are potential therapeutic targets for treating or alleviating the progression of CAVD.

## Materials and methods

### Aortic valve collection

Aortic valve specimens (n=38) were obtained from CAVD patients who underwent aortic valve replacement surgeries at Shanghai Chest Hospital. In this research, 30 specimens intended for immunohistochemistry and tissue staining analyses were fixed in 4% paraformaldehyde, and 8 specimens intended for RT-PCR analysis were stored at −80°C.

The extended methods of tissue delivery and storage were performed according to a previous study[8].

The valves for immunohistochemistry and tissue staining analyses were prepared according to a previous study[8].

The collection and use of samples were approved by the Ethical Committee of Shanghai Chest Hospital. Informed written consent was obtained from each patient prior to inclusion in the study, and all of the investigations conformed to the principles outlined in the Declaration of Helsinki. The patient information is available in **Table 1**.

### Immunohistochemistry

Human valvular tissue specimens were cut into appropriate sizes and subjected to paraffin embedding after fixation in 4% paraformaldehyde, followed by decalcification with 10% ethylenediaminetetraacetic acid (EDTA). Serial sections (4 μm thick) were obtained for immunohistochemistry, Prussian blue, Alizarin red and Masson’s trichrome staining and subjected to antigen retrieval. Endogenous peroxidase was blocked with 3% hydrogen peroxide for 20 min, and nonspecific binding sites were blocked with QuickBlock™ immunostaining blocking reagent (Beyotime, China) for 1 h. The sections were incubated with a primary antibody at 4°C overnight and with a rabbit/mouse HRP-conjugated secondary antibody (Dako, Denmark) at room temperature for 2 h, and proteins were visualized with 3,3-diaminobenzidine (DAB) (Dako, Denmark) for 10 min. All images were captured on an Axio Scope A1 optical microscope (Zeiss, Germany). The primary antibodies used are listed in **the Major Resources Table** in the **Supplementary Materials**.

The positive areas after immunohistochemical staining were quantified and analyzed by Image-Pro Plus 6.0 (Media Cybernetics, USA). All results are presented as the relative integral optical density (IOD) (sum/area sum).

### Perls’ Prussian blue iron staining

A Prussian Blue Iron Stain Kit (Yeasen, China) was used to detect and determine the amounts of iron in the samples in accordance with the manufacturer’s protocol. Iron-containing areas were stained a characteristic blue color by the reagent. All images were captured on an Axio Scope A1 optical microscope (Zeiss, Germany). The positive areas were quantified and analyzed by Image-Pro Plus 6.0 (Media Cybernetics, USA). All results are presented as the relative IOD (sum/area sum).

### Alizarin red staining

To determine the extent of matrix calcification, sections or cultured cells were fixed with 4% paraformaldehyde and then stained with Alizarin Red S (ScienCell, USA) for 30 min in accordance with the manufacturer’s protocol. All photographs were captured on an Axio Scope A1 optical microscope (Zeiss, Germany). All results are presented as the relative IOD (sum/area sum).

### Alkaline phosphatase (ALP) staining

ALP staining was performed using a BCIP/NBT ALP color development kit (Beyotime, China) in accordance with the manufacturer’s instructions. Briefly, cell samples were incubated with ALP-labeled antibodies and washed 3–5 times for 3–5 min each time. Then, each solution was sequentially and washed. After the last wash, a BCIP/NBT dye working solution was added, and the sample was incubated at room temperature for 30 min. All images were captured on an Axio Scope A1 optical microscope (Zeiss, Germany). All results are presented as the relative IOD (sum/area sum).

### Von Kossa staining

Matrix mineralization was assessed using a Von Kossa staining kit (Servicebio, China) in accordance with the manufacturer’s instructions. Briefly, sections or cultured cells were incubated with 1% silver nitrate, placed under UV light for 10 min, and rinsed with distilled water. Unreacted silver was removed by washing with 5% sodium thiosulfate for 5 min. After rinsing with distilled water, the specimens were counterstained with 0.1% Nuclear Fast Red Solution for 5 min. All images were captured on an Axio Scope A1 optical microscope (Zeiss, Germany). All results are presented as the relative IOD (sum/area sum).

### Cell culture and stimulation

Normal human VICs were isolated from healthy sterile valve leaflets obtained from transplant-recipient hearts. The collection and use of human aortic valves were approved by the Ethical Committee of Shanghai Chest Hospital. Immediately after surgical removal, the human aortic valves were immersed in PBS at 4°C and transported to the laboratory. The aortic valve leaflets were digested by collagenase II (Sigma, USA) overnight at 37°C, and endothelial cells were removed using a sterile cotton swab. VICs were placed in high-glucose Dulbecco’s modified Eagle’s medium (Sigma, USA) supplemented with 10% fetal bovine serum (Gibco, USA) and 1% penicillin/streptomycin (Gibco, USA) and seeded onto T-75 flasks. The culture medium was changed every two days. Cells between passages 1 and 3 were used for experiments.

To induce calcification in vitro, VICs were incubated in osteogenic differentiation medium (OM) (ScienCell, USA) for 21 days according to the manufacturer’s instructions. The culture medium was changed every day.

For iron stimulation, VICs were seeded at 1×10^5^ cells/well into 6-well culture plates and stimulated with FeSO_4_ (10 μmol/L) concentrated with iron(II) sulfate heptahydrate (Sigma, USA) in PBS. In these experiments, the culture medium containing FeSO4 was changed every day.

### Real-time quantitative polymerase chain reaction (RT-PCR)

Total RNA was extracted from cultured cells or tissues using an RNA Isolation Kit (Omega Bio-Tek, USA), and the amount and quality were evaluated by spectrophotometry on a Synergy.H1 Microplate Reader (BioTek, USA). cDNA was reverse-transcribed by HiScript III RT SuperMix (Vazyme, China) on a C1000 Touch Thermal Cycler (Bio-Rad, USA), and the amount and quality were evaluated by spectrophotometry on a Synergy.H1 Microplate Reader (BioTek, USA). Real-time PCR was performed using ChamQ SYBR Color qPCR Master Mix (Vazyme, China) on the ViiA7 system (Life Technologies, USA). The relative expression levels of target genes were calculated by the 2^-**ΔΔ**CT^ method, and glyceraldehyde 3-phosphate dehydrogenase (GAPDH) was used as the endogenous control. The sequences of the primers used are included in **the Major Resources Table** in the **Supplementary Materials**.

### Western blot analysis

Total RNA was extracted from cultured cells or tissues using an RNA Isolation Kit (Omega Bio-Tek, USA), and the amount and quality were evaluated by spectrophotometry on a Synergy.H1 Microplate Reader (BioTek, USA). cDNA was reverse-transcribed by HiScript III RT SuperMix (Vazyme, China) on a C1000 Touch Thermal Cycler (Bio-Rad, USA), and the amount and quality were evaluated by spectrophotometry on a Synergy.H1 Microplate Reader (BioTek, USA). Real-time PCR was performed using ChamQ SYBR Color qPCR Master Mix (Vazyme, China) on the ViiA7 system (Life Technologies, USA). The relative expression levels of target genes were calculated by the 2^-**ΔΔ**CT^ method, and glyceraldehyde 3-phosphate dehydrogenase (GAPDH) was used as the endogenous control. The sequences of the primers used are included in **the Major Resources Table** in the **Supplementary Materials**.

### RNA sequencing (RNA-seq)

Total RNA was extracted from VICs, including three groups of osteogenically differentiated VICs and three groups of normal VICs, cultured in T75 flasks using TRIzol reagent (Life Technologies, USA) according to the manufacturer’s recommended protocol. The quantity and purity of the RNA were measured using a NanoPhotometer^®^ spectrophotometer (Implen, German), and the RNA integrity was assessed using an Agilent Bioanalyzer 2100 (Agilent Technologies, USA).

Prior to library preparation, ribosomal RNAs (rRNAs) were removed from the total RNA. The rRNA-depleted RNAs were then fragmented to obtain short 300 bp fragments, converted into cDNA, connected to adapters, sorted and amplified by PCR. Using the NextSeq 500/550 High Output Kit v2 (150 cycles, Illumina, USA), 75 bases in both the forward and reverse directions of the library were sequenced on an Illumina NextSeq machine.

For bioinformatics data analysis, the raw sequencing data were analyzed by FAST-QC (http://www.bioinformatics.babraham.ac.uk/projects/fastqc/) to filter the low-quality and adaptor sequences. Then, the clean reads were mapped to the human reference genome by HISAT2 (version 2.1.0; https://daehwankimlab.github.io/hisat2/). The mRNA levels in VICs were quantified as counts and fragments per kilobase of transcript per million mapped reads (FPKM), and the differentially expressed genes (DEGs) with a false discovery rate (FDR) < 0.05 and a log2 fold change >1 were identified by the DE-Seq2 algorithm. The functions of the main DEGs were determined by Gene Ontology (GO) analysis[10], and Kyoto Encyclopedia of Genes and Genomes (KEGG) (http://www.genome.jp/Kegg/) analysis was performed to identify the enriched pathways among the DEGs. Fisher’s exact test was used to identify the significantly enriched pathways. The P values were adjusted using the BH FDR algorithm, and the significance threshold for the GO and KEGG analyses was an FDR < 0.05.

### Plasmids

For plasmid transfection, VICs were seeded at 5,000 cells/well into 96-well culture plates, transfected with either an empty vector (Solarbio, China) or a Slc7a11 cDNA ORF Clone (NM_014331.3) (SinoBiological, China) using Lipofectamine 3000 Transfection Reagent (Invitrogen, USA) and cultured for 24 h in minimal medium comprising DMEM and 5% FBS without antibiotics. Then, the VICs were harvested for transient transfection analysis, and successful overexpression was confirmed by RT-PCR.

### Lentiviral transfection

Lentiviruses carrying Slc7a11 shRNAs were designed and produced by Genomeditech (China). VICs were seeded at 1×10^5^ cells/well into 12-well culture plates, and the appropriate lentiviruses were then added to each well (MOI: 50~80). The medium in each well was replaced after 2 days, and the cells were screened after two more days using antibiotics recommended by the company. Then, the VICs were harvested for stable transfection analysis, and three different shRNAs were used to exclude off-target effects. These shRNA sequences are listed in **the Major Resources Table** in the **Supplementary Materials**. Successful knockdown was confirmed by RT-PCR (**Supplementary Figure S1**). After measuring the transfection efficiency, shRNA#1 was chosen for further studies.

### Transmission electron microscopy

After the indicated treatment, VICs grown on a 100-mm dish were quickly harvested and immediately fixed in 3% phosphate-glutaraldehyde in 0.1 M PBS at room temperature for 1 h, post-fixed in 1% OsO4 in 0.1 M PBS at room temperature for 1 h, dehydrated using a graded series of ethanol, embedded with epoxy resin and sectioned. The sections were then examined under a Tecnai 10 (100 kV) transmission electron microscope (FEI), and at least three random views were recorded for each sample.

### ROS, lipid peroxidation, NADPH, GSH, total iron and ferrous ion measurements

ROS were measured using fluorescence-activated cell sorting with an ROS Assay Kit (Beyotime, China). Lipid peroxidation, NADPH and glutathione (GSH) were measured using a colorimetric Lipid Peroxidation Assay Kit (Abcam, UK), NADPH Assay Kit (Abcam, UK) and GSH Assay Kit (Abcam, UK), respectively. The total iron and ferrous ion contents were measured using an iron assay (Abcam, UK). All experiments were performed according to the manufacturers’ instructions.

### Statistical analysis

Statistical analyses were performed with GraphPad Prism (version 5.01). The data were analyzed with Student’s t test, one-way ANOVA, and the Dunnett test, and p<0.05 indicated statistical significance.

### Data availability statement

The RNA-seq data are available in GenBank at NCBI.

## Results

### Ferroptosis occurs in IH areas

By analyzing different areas of calcific valve specimens with IH (n=30), we positively identified iron in each area of the calcific valves. Unsurprisingly, the positively iron-stained areas were largest in the IH areas (**Figure 1A,C**). To evaluate the level of 4-hydroxynonenal (4-HNE) (a key feature of ferroptosis), immunohistochemical staining showed that 4-HNE was highly expressed in the IH areas (**Figure 1D**), indicating aberrant lipid peroxidation in the IH areas. Moreover, we assessed the contents of iron, NADPH, and GSH and lipid peroxidation in different areas of aortic valve specimens. In the IH areas, iron mass accumulation was found, the NADPH content decreased (**Figure 1E,F**), the lipid peroxidation increased (**Figure 1G**), and the GSH content decreased (**Figure 1H**). Moreover, using flow cytometry to detect the ROS content in VICs from different areas, we found that VICs from IH areas had high ROS contents (**Figure 1I**). These data suggested that ferroptosis may be associated with IH in the pathological progression of CAVD.

**Figure 1.**
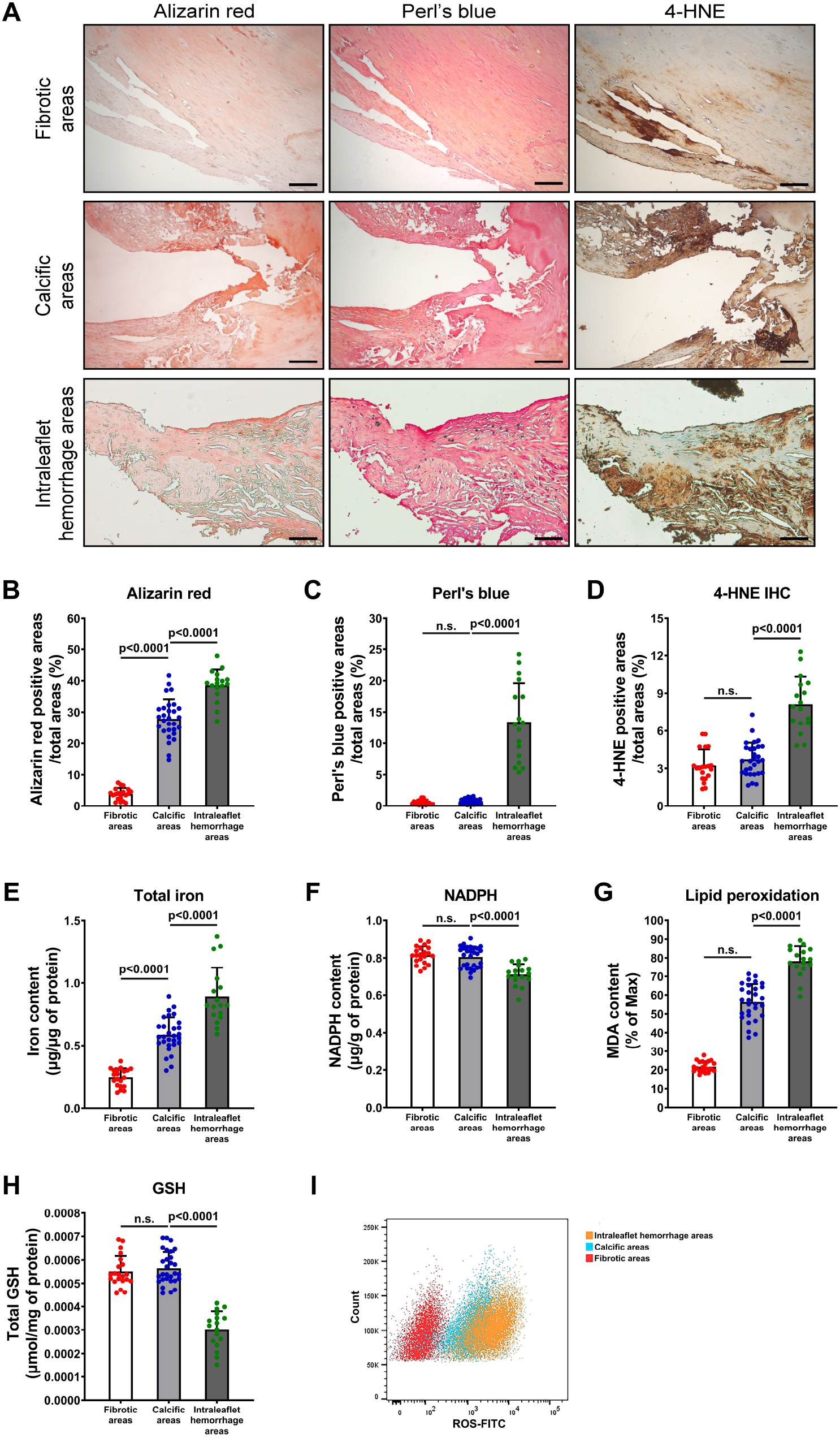
Ferroptosis occurs in intraleaflet hemorrhage areas. **A.** Serial cross-sections of human calcific aortic valves with IH (n=30) were stained with Alizarin red, Perls’ blue or 4-HNE immunochemistry. Scale bar, 100 μm. **B-D.** The areas positively stained with Alizarin red (**B**), Perls’ blue (**C**) and 4-HNE (**D**) were quantified. **E-I.** The total iron content (**E**), NADPH content (**F**), lipid peroxidation (**G**), GSH content (**H**) and ROS content (**I**) in fibrotic, calcific and IH areas. Measured data are presented as the mean ± SD.

### Slc7a11 expression is downregulated in OM-treated VICs and calcified aortic valves

To elucidate an unbiased expression profile of the genes involved in VIC osteogenic differentiation, RNA-seq was used to compare normal VICs and OM-treated VICs to identify potential gene candidates. The genes associated with calcification, fibrosis and inflammation were truly upregulated in OM-treated VICs compared with normal VIC controls. However, Slc7a11 was significantly downregulated in OM-treated VICs (**Figure 2A**), which attracted our attention because Slc7a11 plays an important role in ferroptosis[11,12]. Interestingly, GO biological process and pathway analyses also suggested the enrichment of negative oxidative stress regulation, cellular iron ion homeostasis and GSH metabolism in OM-treated VICs (**Supplementary Figure S2A,B**). Next, we verified the mRNA levels of iron metabolism-related genes in fibrotic areas (n=8), calcific areas (n=8) and IH areas (n=8). Slc7a11 was significantly differentially expressed in calcific and IH areas (**Figure 2B**). We reconfirmed the expression of Slc7a11 in VICs after treatment with OM and DMEM and found that the mRNA and protein levels of Slc7a11 were decreased in OM-treated VICs in a time-dependent manner (**Figure 2C,D; Supplementary Figure S3**). As the expression and role of Slc7a11 in calcific aortic valves have not been previously documented, we hypothesized that Slc7a11 may participate in the mechanism of CAVD.

**Figure 2.**
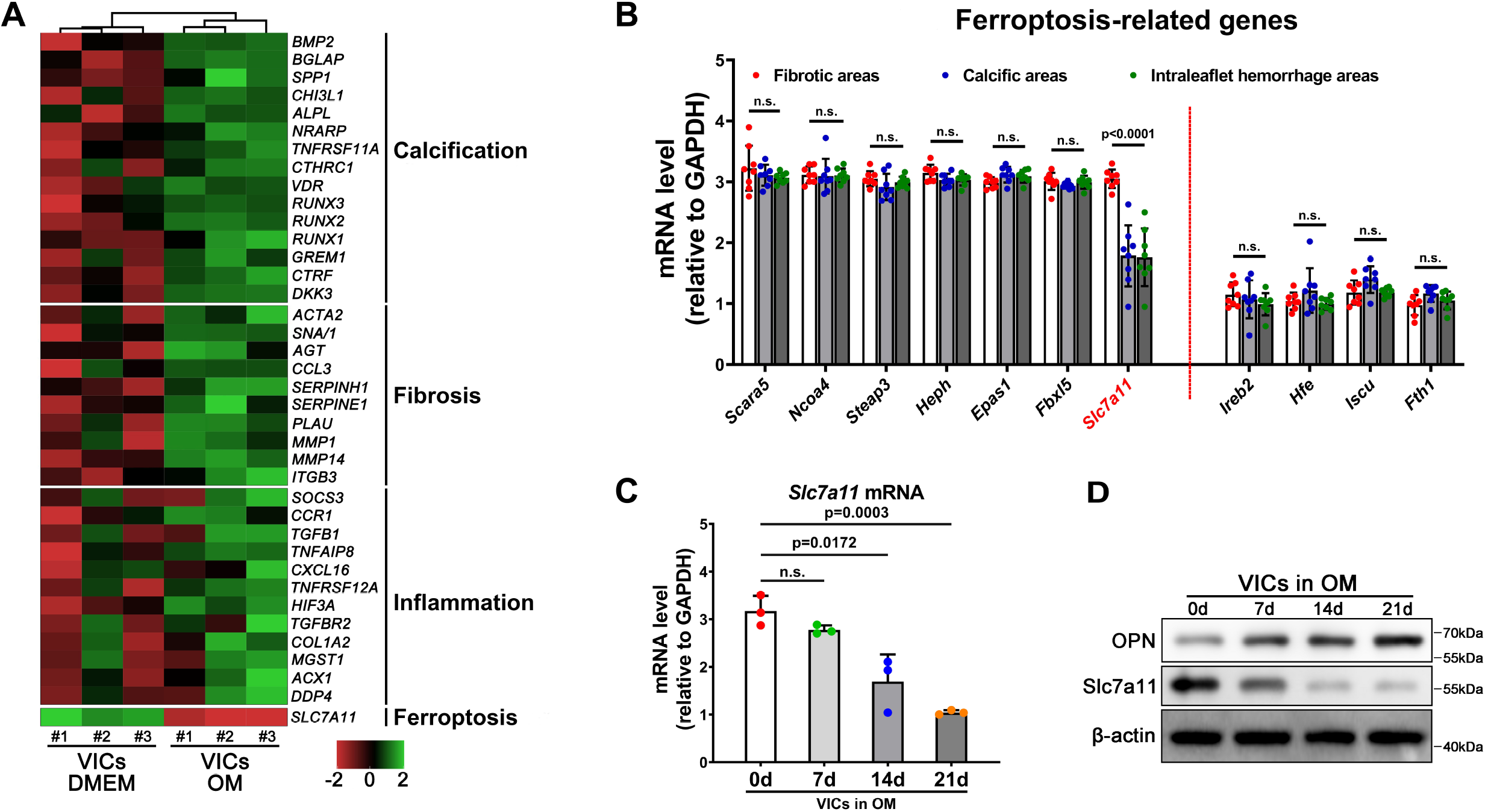
Slc7a11 expression is downregulated in OM-treated VICs and calcified aortic valves. **A.** Gene expression profiles of normal VICs and OM-treated VICs. Up- and downregulated transcripts are shown in red and green, respectively. **B.** The mRNA expression levels of ferroptosis-related genes in fibrotic areas (n=8), calcific areas (n=8) and IH areas (n=8) were evaluated by RT-PCR. GAPDH was used for normalization. **C-D.** The mRNA (**C**) and protein (**D**) expression levels of Slc7a11 in normal VICs and OM-treated VICs were evaluated by RT-PCR and western blot analyses. GAPDH and β-actin was used for normalization. Each experiment was performed three times, and measured data are presented as the mean ± SD.

### Low Slc7a11 expression is associated with calcification in CAVD

Sections of surgical aortic valve specimens (n=30) from different areas were stained with Alizarin red or Von Kossa stain to assess the degree of calcification, stained with Perls’ Prussian blue to detect iron, and incubated with antibodies against Slc7a11 to detect Slc7a11 expression. As shown in **Figure 3A**, compared with that in fibrotic areas, Slc7a11 expression in calcific and IH areas was lower. The sections positively stained with Alizarin red, Von Kossa stain, Prussian blue and anti-Slc7a11 antibodies were quantified using computer-aided planimetry and are expressed as a percentage of the total surface area (**Figure 3B-E**). These quantified results were consistent with the immunohistochemistry results. Taken together, the results indicated that low Slc7a11 expression may be associated with calcific aortic valves.

**Figure 3.**
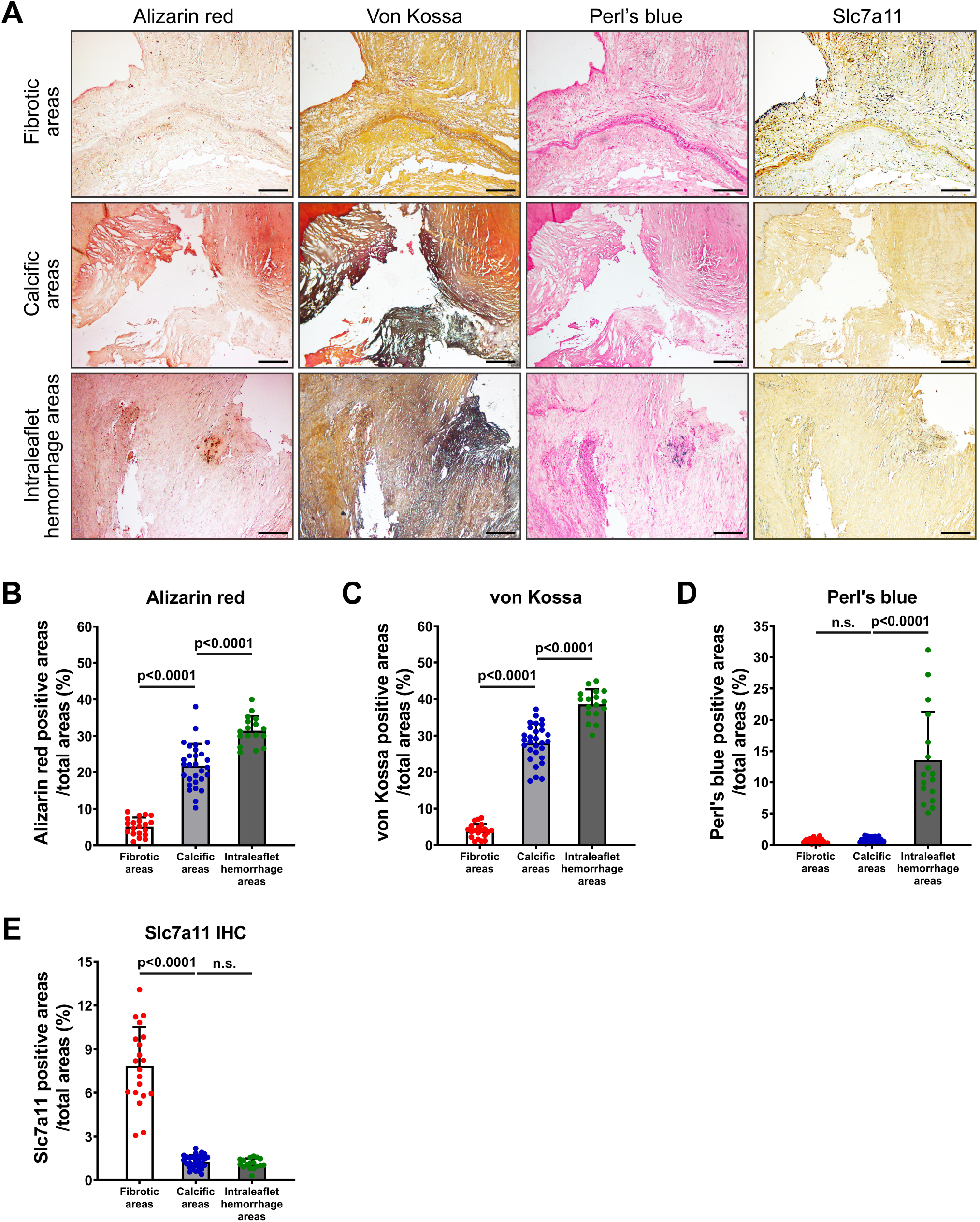
Low Slc7a11 expression is associated with calcification in CAVD. **A.** Serial cross-sections of human calcific aortic valves with IH (n=30) were stained with Alizarin red, Von Kossa, Perls’ blue or Slc7a11 immunochemistry. Scale bar, 100 μm. **B-E.** The areas positively stained with Alizarin red (**B**), Von Kossa (**C**), Perls’ blue (**D**) and 4-HNE (**E**) were quantified. Measured data are presented as the mean ± SD.

### Long-term iron overload does not promote VIC calcification under osteogenic conditions

To determine the effect of iron on aortic valve calcification, VICs were cultivated in OM only or with added iron. After 21 days of iron stimulation, the total iron and ferrous ion contents in VICs were increased by approximately fourfold (**Figure 4A,B**), indicating that iron was absorbed by the VICs under osteogenic conditions. OPN, osteocalcin (OCN), Runx2 and Osterix were used as markers of osteogenic differentiation. As shown in **Figure 4C**, compared with that in VICs cultured in OM, the mRNA expression levels of OPN did not change. However, the OCN, Runx2 and Osterix mRNA levels were enhanced in VICs with long-term iron overload cultured under osteogenic conditions (**Figure 4D-F**). We also tested the protein levels of osteogenic markers by western blot analysis and found that the protein expression of these osteogenic markers did not increase (**Figure 4G**), which indicated that iron stimulation alone did not significantly promote the osteogenic differentiation of VICs at the protein level. Additionally, the osteoinductive effect of iron on VICs was confirmed by Alizarin red, ALP and Von Kossa staining. Compared with that in cells cultured in OM, calcium deposition in cells cultured with iron in OM was not obviously increased (**Figure 4G**). Quantification of the positive Alizarin red, ALP and Von Kossa staining areas by computer-aided planimetry also supported that iron stimulation alone did not promote VIC calcification (**Figure 4H-J**).

**Figure 4.**
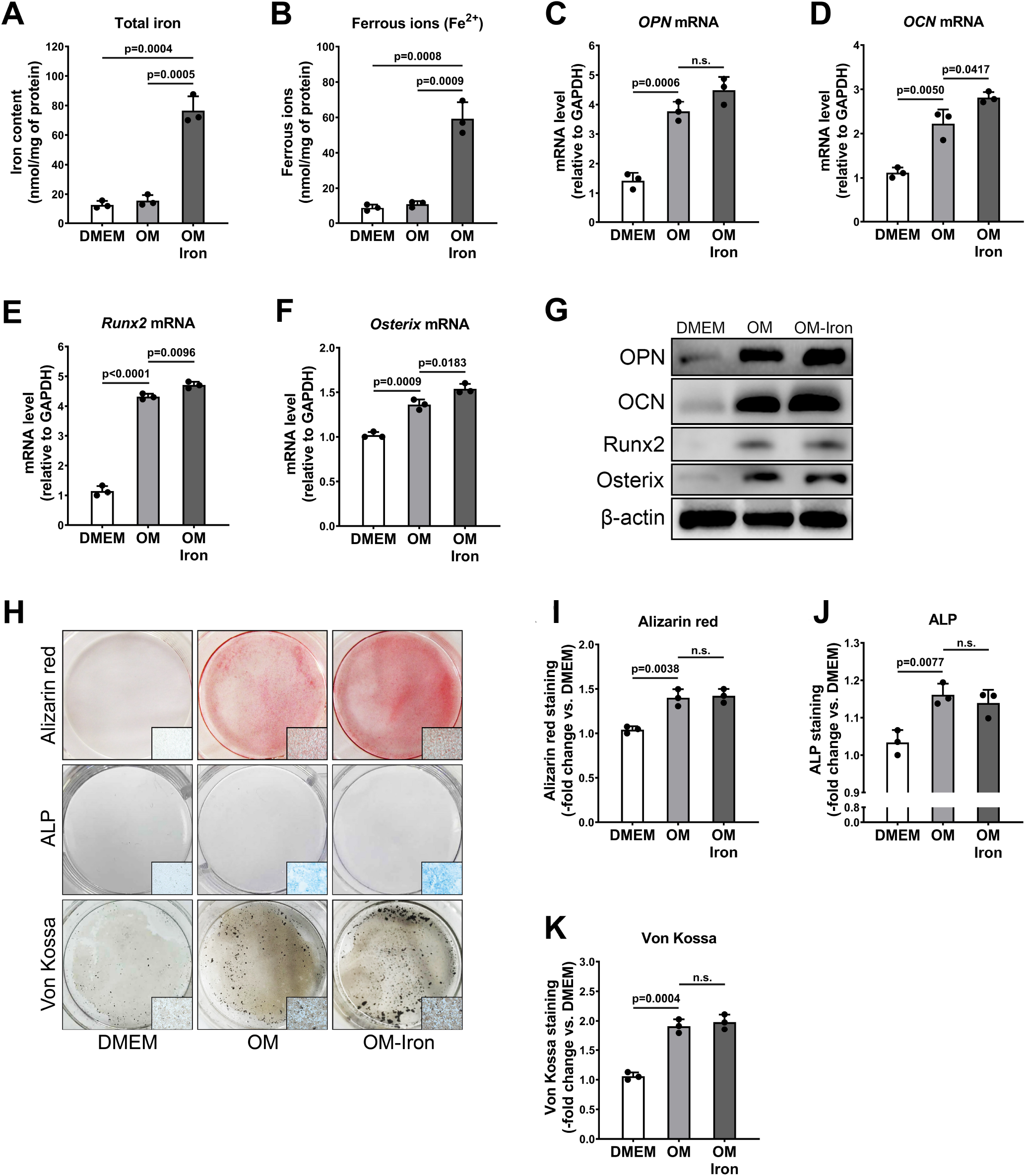
Long-term iron overload does not promote VICs calcification under osteogenic conditions. **A-B.** The contents of iron (**A**) and ferrous ion (**B**) in VICs. **C-F.** The mRNA expression levels of OPN (**C**), OCN (**D**), Runx2 (**E**) and Osterix (**F**) in VICs after treatment with OM and iron were evaluated by RT-PCR. GAPDH was used for normalization. **G-J.** The calcific deposition of VICs after treatment with OM and iron was observed by alizarin red staining, ALP staining and Von Kossa staining (**G**). The positively stained areas were quantified (**H-J**). Each experiment was performed three times, and measured data are presented as the mean ± SD.

### Slc7a11 deficiency aggravates iron overload-induced ferroptosis in vitro

As Slc7a11 is expressed at low levels in calcific aortic valves, we next assessed whether Slc7a11 also participates in the mechanism of VIC calcification under iron overload. First, we successfully overexpressed Slc7a11 (ov-Slc7a11) and knocked down Slc7a11 (si-Slc7a11) in VICs (**Figure 5A,B**). Then, we explored whether Slc7a11 regulated iron transport in VICs. Compared with those in normal VICs treated with iron, the total iron and ferrous ion contents in VICs with Slc7a11 overexpression or knockdown were not different (**Figure 5C,D**), which indicated that Slc7a11 did not regulate iron transport. We also measured the mRNA expression of Slc7a11 in VICs in DMEM after stimulation with iron and found that its expression was not changed in iron-overloaded VICs, which suggested that osteogenic conditions, rather than iron, regulated the expression of Slc7a11 in VICs (**Supplementary Figure S4**). Then, we assessed ferroptosis in VICs by measuring the ROS, NADPH, and GSH contents and lipid peroxidation. Treatment with iron significantly increased the lipid peroxidation content (**Figure 5E**), decreased the NADPH content (**Figure 5F**), and increased the ROS content (**Figure 5G**) in VICs. However, overexpression of Slc7a11 in VICs effectively inhibited the level of ferroptosis induced by iron overload (**Figure 5E-G**). In contrast, knockdown of Slc7a11 aggravated iron overload-induced ferroptosis in VICs (**Figure 5E-G**). Changes in mitochondrial morphology are also characteristic of iron overload-induced ferroptosis[13]. Transmission electron microscopy revealed that compared with normal VICs, si-Slc7a11 VICs grown under osteogenic conditions exhibited shrunken mitochondria with an increased membrane density; however, overexpression of Slc7a11 inhibited mitochondrial shrinkage and rupture (**Figure 5H, I**). These results suggested that Slc7a11 deficiency aggravates iron overload-induced ferroptosis in VICs.

**Figure 5.**
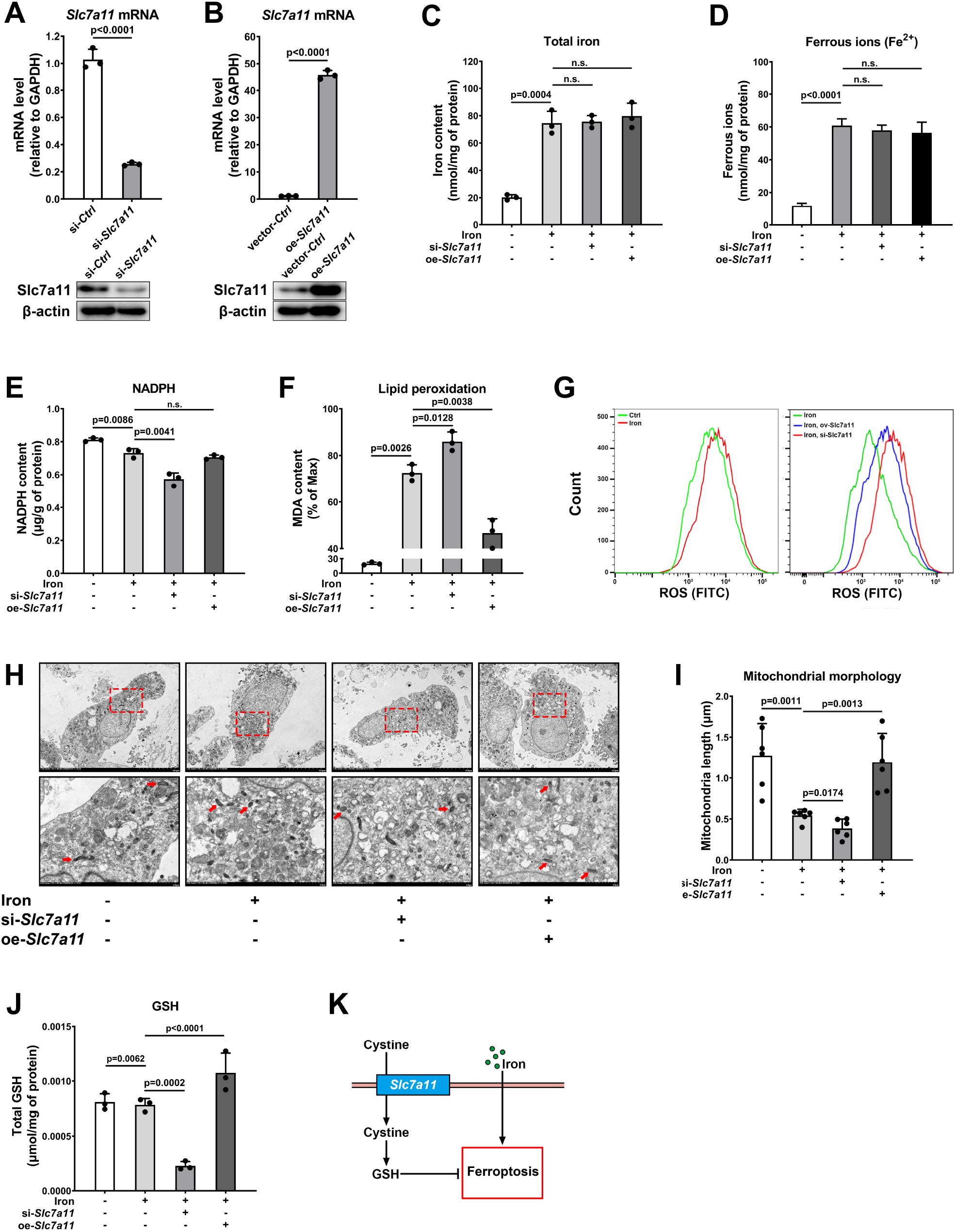
Slc7a11 deficiency aggravates iron overload-induced ferroptosis in vitro. **A-B.** Slc7a11 was knocked down (**A**) and overexpressed (**B**) in VICs. **C-D.** The contents of iron (**C**) and ferrous iron (**D**) in normal VICs, si-Slc7a11 VICs and ov-Slc7a11 VICs after treatment with iron. **E-G.** The NADPH content (**E**), lipid peroxidation (**F**) and ROS content (**G**) in normal VICs, si-Slc7a11 VICs and ov-Slc7a11 VICs after treatment with iron. **H-I.** Normal VICs, si-Slc7a11 VICs and ov-Slc7a11 VICs were treated with iron and then examined using transmission electron microscopy (**H**). The mitochondrial length (along the long axis) was measured and summarized (**I**). **J.** Total GSH in normal VICs, si-Slc7a11 VICs and ov-Slc7a11 VICs after treatment with iron. **K.** Schematic diagram depicting the key function of Slc7a11 in GSH biosynthesis and iron overload-induced oxidative stress. Each experiment was performed three times, and measured data are presented as the mean ± SD.

GSH prevents cell damage caused by ROS, such as free radicals and peroxides. We next measured the GSH content and found that Slc7a11 knockdown decreased the total GSH in VICs (**Figure 5J**), which indicated that Slc7a11 regulated iron overload-induced ferroptosis via GSH. The in vitro data shown in **Figure 5K** indicate that Slc7a11 deficiency decreases the synthesis of GSH and aggravates ferroptosis induced by iron overload in VICs. Briefly, Slc7a11 deficiency decreases the ability of VICs to resist ferroptosis induced by iron overload.

### Iron promotes Slc7a11-deficient VIC osteogenic differentiation by aggravating ferroptosis in vitro

We next explored whether the loss of Slc7a11 could facilitate VIC calcification after iron overload in vitro, as we observed in the surgical specimens. In this experiment, Slc7a11 was knocked down in VICs, and all VICs were cultured with iron in OM. Some groups of VICs were treated with the iron chelator deferoxamine (DFO), the ROS inhibitor acetylcysteine (Acet) or the specific ferroptosis inhibitor ferrostatin-1 (Fer-1). The iron and ferrous ion contents in each group were not altered by treatment with DFO, which suggested that these inhibitors did not affect iron transport in VICs (**Figure 6A, B**). Iron overload-induced calcification was confirmed by the osteogenic differentiation markers OPN, OCN, Runx2 and Osterix. Compared with those in normal VICs treated with iron, the mRNA and protein levels of OPN, OCN, Runx2 and Osterix in VICs with Slc7a11 knockdown were downregulated under osteogenic conditions (**Figure 6C-G**). In contrast, Acet, Fer-1 and DFO significantly reduced iron overload-induced calcification in si-Slc7a11 VICs (**Figure 6C-G**). The osteoinductive effect of iron on VICs was also confirmed by Alizarin red, ALP and Von Kossa staining. Slc7a11 deficiency obviously increased calcium deposition in VICs grown in OM after treatment with iron, and Acet, Fer-1 and DFO significantly rescued iron overload-induced calcification (**Figure 6H**). The quantification of positive Alizarin red, ALP and Von Kossa staining areas by computer-aided planimetry also supported that iron stimulation alone did not promote VIC calcification (**Figure 6I-K**). These in vitro data suggested that the loss of Slc7a11 facilitates iron overload-induced calcification by aggravating ferroptosis stress in VICs.

**Figure 6.**
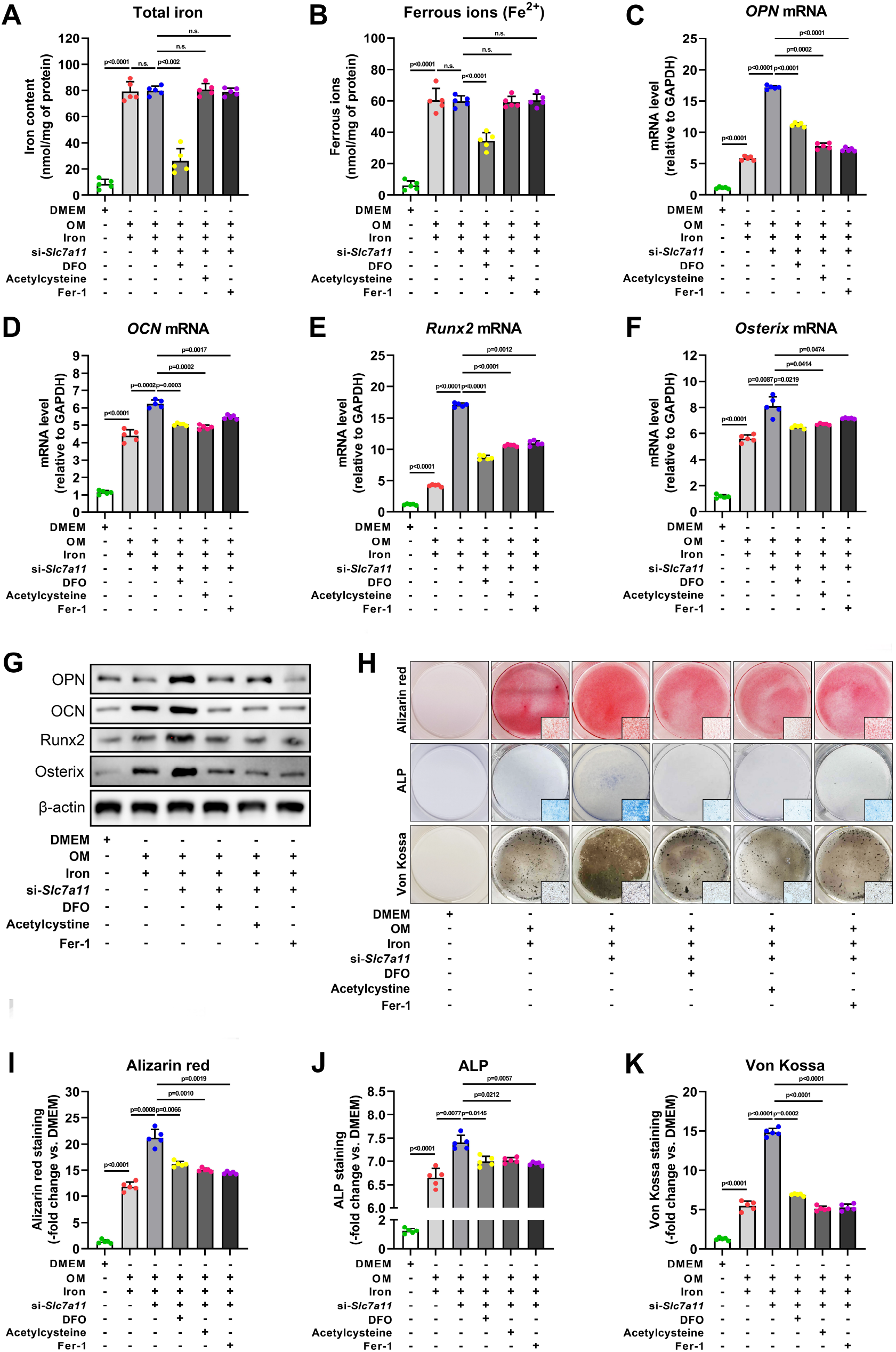
Iron promotes Slc7a11-deficient valvular interstitial cell osteogenic differentiation by aggravating ferroptosis in vitro. **A-B.** The contents of iron (**A**) and ferrous ions (**B**) in VICs after treatment with DFO, Aect and Fer-1 in OM and iron stimulation. **C-F.** The mRNA expression levels of OPN (**C**), OCN (**D**), Runx2 (**E**) and Osterix (**F**) in VICs after treatment with DFO, Aect and r-GSH in OM with iron stimulation were evaluated by RT-PCR. GAPDH was used for normalization. G. The protein expression levels of OPN, OCN, Runx2 and Osterix in VICs after treatment with DFO, Aect and r-GSH in OM with iron stimulation were measured by western blot analysis. β-actin was used for normalization. **H-K.** The calcific deposition of VICs after treatment with DFO, Aect and r-GSH in OM and iron stimulation was observed by Alizarin red staining, ALP staining and Von Kossa staining (**H**).

## Discussion

CAVD is characterized by the abnormal long-term accumulation of calcium-rich nodules on the aortic valve cusp, leading to thickening (termed sclerosis), limited movement, and stenosis (left ventricular outflow obstruction)[14]. VICs are the major cell type of aortic valves, and calcium salts in diseased aortic valves are mainly derived from osteogenically differentiated VICs. Although inflammation, lipid accumulation, matrix remodeling, angiogenesis, the renin-angiotensin system and some other mechanisms and pathways of VICs osteogenic differentiation have been described, there are still many mechanisms that initiate or accelerate the calcification of aortic valve leaflets that need to be explored[1,13,15].

IH may participate in the process of CAVD[5,7]. In IH areas, red blood cell (RBC) metabolism of macrophages and neovascularization may cause the accumulation of iron. Marion M et al. reported that VICs may acquire an osteoblastic phenotype by contacting senescent RBCs, which aggravates calcium deposition in human aortic valves[5]. Additionally, Laguna-Fernandez A et al. found that iron can be taken up by VICs in a proinflammatory environment and actively contribute to VIC proliferation and extracellular matrix remodeling[7]. Although these studies mainly focused on the histological analysis of calcific aortic valve leaflets and suggested that iron accumulation is associated with calcification in CAVD[5,7], it is still necessary to deeply explore the mechanism of iron accumulation and calcium deposition through in vivo and in vitro experiments.

Iron homeostasis is essential for the proper functioning of all mammalian cell types, including VICs. As cardiac tissue is highly vulnerable to the accumulation of free iron[16], abnormal iron-induced toxicity (ferroptosis) plays a role in the pathogenesis of many cardiovascular diseases[17]. Xuexian F et al. found that ferritin plays a major role in protecting against cardiac ferroptosis and subsequent heart failure[18]. Tao B et al. reported that inhibition of ferroptosis could alleviate atherosclerosis by attenuating lipid peroxidation and endothelial dysfunction in aortic endothelial cells[19]. Although research on ferroptosis in cardiovascular diseases is widespread, specific research on its mechanism in CAVD is still lacking. In the present study, iron was consistently visualized and accumulated gradually throughout CAVD progression. The tissue analysis results strongly suggest that iron is associated with calcification in CAVD, as previously reported.

Slc7a11 is a major component of the glutamate/cystine antiporter system xc, which regulates the downstream synthesis of GSH, which is a key antioxidant that scavenges lipid peroxides[20]. Many studies have demonstrated that Slc7a11 plays an important role in antioxidative stress[21–23] and maintains normal iron metabolism[24]. Reduced Slc7a11 expression increases oxidative stress and plays an important role in the mechanism of cardiovascular calcification[25,26].

In this research, two novel and important notions were proposed. First, we first reported that ferroptosis is associated with the progression of CAVD. Second, this is the first VIC cytology experiment to clarify the mechanism by which IH aggravates valve calcification. Considering all of these results, we report a potential mechanism by which iron promotes Slc7a11 deficiency in VIC osteogenic differentiation. From a clinical perspective, our results suggest that inhibiting ferroptosis and IH may be a new therapeutic strategy for alleviating the progression of CAVD, especially in patients who have calcified aortic valves without severe stenosis.

### Conclusion

In conclusion, iron promotes Slc7a11-deficient VIC osteogenic differentiation by aggravating ferroptosis in vitro, thereby accelerating the progression of aortic valve calcification.

### Study limitations

This research involved only histological and cytological experiments and lacked animal experiments because there are no mature animal models of IH in calcific aortic valves; thus, it is difficult to simulate localized iron overload in an animal model. Our research focus on factors that accelerate the progression of CAVD, not the factos that initiate the progression of CAVD. We also analyzed the levels of iron overload and ferroptosis in an ApoE^-/-^ mouse model (fed a Western diet for 4 months) and did not find an obvious change.

Another limitation is that we did not further research the upstream region of Slc7a11. It will be valuable to characterize the regulation of Slc7a11 expression under osteogenic conditions in our future research. Moreover, the mechanism by which oxidative stress promotes calcification has already been reported many times; therefore, we did not repeat these downstream studies of ferroptosis.

## Acknowledgments

None.

